# Integrated single-cell and spatial transcriptomic analysis of T cell exhaustion and immunometabolic remodeling in HPV-positive oropharyngeal squamous cell carcinoma

**DOI:** 10.64898/2026.03.11.710950

**Authors:** Kexin Wang, Nian Deng, Yueqin Tao, Xiaochen Yang, Mujie Yuan, Longhua Gao, Shuai Jiang, Wei Shang, Jing Deng, Lin Wang

## Abstract

HPV-positive oropharyngeal squamous cell carcinoma (OPSCC) harbors dense lymphocytic infiltration yet responds poorly to immune checkpoint blockade (ICB), a paradox whose mechanistic basis remains unresolved. Here we constructed an integrated single-cell and spatial transcriptomic atlas of HPV⁺ OPSCC (23,119 cells, 24 cell types), complemented by CRISPR-Cas9 functional validation in two HPV⁺ cell lines and multiplex immunofluorescence across independent cases. Pseudotime trajectory analysis placed TOX⁺CD8⁺ T cells, rather than canonically defined exhausted cells, at the terminal exhaustion endpoint, and uncovered a previously uncharacterized IEG⁺CD8⁺ effector memory population at a fate-decision juncture between activation and irreversible exhaustion. Actively cycling Tregs with broad co-stimulatory and co-inhibitory receptor expression underwent intratumoral immunosuppressive differentiation. Compartmentalized LDHA/MCT4 expression in cancer cells versus LDHB/MCT1 in T cells established a directional lactate flux model, and spatial deconvolution confirmed peripheral immune exclusion. We identified a KMT2D-KLF7-PD-L1 regulatory axis: KMT2D sustains KLF7 transcription through H3K4me1-dependent enhancer activation; KLF7 concurrently drives neural crest differentiation and upregulates PD-L1, thereby linking epigenetic remodeling to both neural programming and immune checkpoint expression. These findings define converging multi-scale mechanisms underlying ICB resistance in HPV⁺ OPSCC and nominate the KMT2D-KLF7-PD-L1 axis as a combinatorial therapeutic target.

## Introduction

Oropharyngeal squamous cell carcinoma (OPSCC) driven by human papillomavirus (HPV) infection has become one of the most rapidly increasing malignancies in high-income countries, currently accounting for approximately 70% of OPSCC cases in the United States and over 50% in the United Kingdom.^1,2^ The eighth edition of the UICC/AJCC staging system formally separates HPV-positive from HPV-negative OPSCC, reflecting the substantially better prognosis of HPV-driven disease.^3^ HPV⁺ OPSCC retains wild-type TP53, is driven by viral oncoprotein (E6/E7)-mediated cell cycle dysregulation, and harbors dense lymphocytic infiltrates with higher CD8⁺ T cell density, elevated PD-L1 expression and more active interferon signaling than its HPV-negative counterpart.^4–7^ Yet in the CheckMate-141 and KEYNOTE-048 trials, objective response rates to anti-PD-1 monotherapy remained below 20% even among PD-L1-positive patients,^8,9^ indicating that functional heterogeneity within the immune microenvironment, rather than total immune cell density, governs therapeutic responsiveness.^10^ Deciphering why an immunogenic tumor resists immunotherapy demands dissection of cell states, spatial organization, metabolic context and tumor-intrinsic programs that collectively shape the local immune landscape.

The cellular basis of this resistance likely originates with T cell dysfunction. CD8⁺ T cells differentiate along a continuum from cytotoxic effectors to terminally exhausted populations co-expressing PD-1, TIM-3, LAG-3 and the transcription factor TOX, which epigenetically enforces a dysfunctional state refractory to ICB-mediated reinvigoration.^11–14^ Whether the exhaustion trajectory in HPV⁺ OPSCC follows a disease-specific path with distinct branching points is unknown. The immediate early gene (IEG) transcription factors FOS and JUN, rapidly induced upon TCR engagement, are master regulators of T cell activation,^15^ and c-Jun overexpression confers exhaustion resistance in CAR-T cells,^16^ but whether IEG-driven programs define a discrete fate-decision population between activation and terminal exhaustion has not been explored. In the CD4⁺ compartment, Tregs marked by FOXP3, CTLA-4, OX40 and GITR suppress antitumor immunity and can paradoxically expand after anti-PD-1 therapy,^17,18^ yet their intratumoral differentiation dynamics in HPV⁺ OPSCC remain uncharacterized. A high-resolution map of both effector and suppressive T cell trajectories is therefore needed to pinpoint where therapeutic intervention could redirect immune cell fate.

T cell dysfunction does not occur in a metabolic vacuum. Cancer cells elevate LDHA and MCT4 (SLC16A3) to drive lactate efflux,^19,20^ creating a lactate-rich milieu that suppresses CD8⁺ T cell cytotoxicity while supporting Treg survival.^21,22^ In HNSCC, spatial transcriptomics has revealed metabolic gene expression patterns that define immunosuppressive niches,^23^ but the spatial relationship between metabolic compartmentalization and T cell exhaustion gradients in HPV⁺ OPSCC has not been examined. Recent single-cell studies have characterized HNSCC microenvironment heterogeneity,^24–28^ and computational tools such as CellChat enable systematic inference of intercellular communication.^29,30^ Despite these advances, no study has integrated single-cell transcriptomics with spatial profiling to simultaneously map T cell trajectories, metabolic compartmentalization and immune topology in HPV⁺ OPSCC, leaving unanswered whether the immune-excluded spatial phenotype^31^ mechanistically underlies ICB resistance.

Tumor-intrinsic molecular programs may also actively shape the immunosuppressive microenvironment. KMT2D (MLL4), a histone methyltransferase mutated in ∼17% of HNSCC,^32,33^ catalyzes enhancer H3K4 monomethylation required for gene activation,^34^ and its loss deregulates interferon signaling and immune microenvironment composition.^35,36^ Computational analyses have further uncovered synthetic lethal interactions between KMT2D loss and DNA repair genes,^37^ suggesting that KMT2D status may simultaneously influence immune regulation and create therapeutic vulnerabilities. Whether KMT2D-dependent enhancer activity regulates specific downstream transcription factors linking epigenetic remodeling to immune evasion in HPV⁺ OPSCC has not been investigated. One candidate mediator is KLF7, a transcription factor that drives neural crest differentiation in embryonic neuroblasts.^38^ Cancer-associated neurogenesis promotes tumor progression and immune evasion,^39^ perineural invasion is an adverse prognostic feature in HNSCC,^40^ and Schwann cells contribute to perineural invasion through ECM remodeling.^41^ Yet whether tumor cells themselves acquire neural crest-like transcriptional programs linked to immune checkpoint expression remains entirely unexplored.

To address these convergent gaps, we performed integrated scRNA-seq and Visium spatial transcriptomics on HPV⁺ OPSCC, complemented by CRISPR-Cas9 KMT2D knockout, KLF7 overexpression and multiplex immunofluorescence across independent cases. We constructed a 24-cell-type atlas, traced branched CD8⁺ and CD4⁺ T cell trajectories, mapped intercellular signaling networks and immunometabolic compartmentalization, and uncovered a KMT2D-KLF7-PD-L1 regulatory axis linking enhancer-mediated epigenetic remodeling to DNA repair vulnerability, neural crest differentiation and immune checkpoint expression.

## Results

### Single-cell atlas reveals a T cell-dominant microenvironment in HPV^+^ OPSCC

scRNA-seq yielded 23,119 high-quality cells after quality control and doublet removal. Unsupervised clustering identified 24 distinct cell types (Fig. 1a). T lymphocytes constituted the predominant population, consistent with the immunologically “hot” phenotype of HPV-driven oropharyngeal cancers. Six CD8⁺ T cell functional states were resolved: CD8_T_cytotoxic, CD8_T_effector_memory, CD8_T_exhausted, IEG⁺CD8_Tem, TOX⁺CD8_T and CD8_T_cycling. The CD4⁺ compartment comprised CD4_T_naive/centr_mem, CD4_T_exhausted, Treg and Treg_Cycling. The atlas additionally captured B cells, plasma cells, monocytes/macrophages, conventional dendritic cells (cDC), plasmacytoid dendritic cells (pDC), mature regulatory dendritic cells (mreg_DC), NKT cells, a heat shock protein-associated stressed T cell population (HSP_stressed_T), cancer cells, endothelial cells and stromal cells (Fig. 1b). Cell cycle analysis confirmed that cycling populations were confined mainly to CD8_T_cycling, Treg_Cycling and a fraction of cancer cells (Fig. 1c). The coexistence of six CD8⁺ T cell functional states and actively cycling Tregs within a single tumor points to immune heterogeneity far exceeding what bulk transcriptomic studies have suggested, implicating the composition and functional states of infiltrating populations-rather than insufficient immune cell recruitment-as the basis for ICB failure in this disease.

**Fig. 1.**
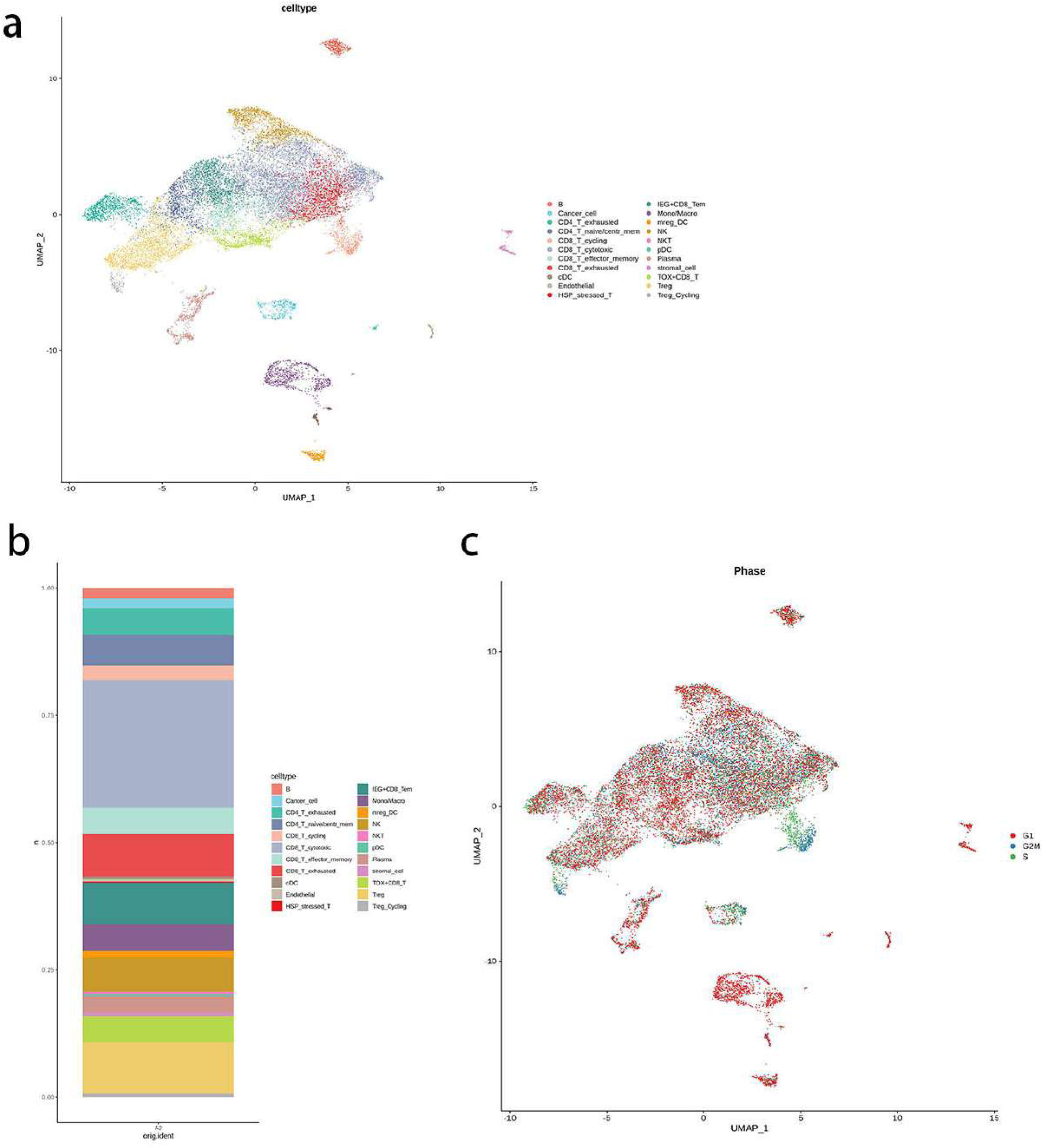
Single-cell transcriptomic atlas of HPV+ OPSCC. **a** UMAP visualization of 23,119 cells annotated into 24 cell types. **b** Stacked bar plot showing the proportion of each cell type. **c** UMAP colored by cell cycle phase (G1, G2M, S), demonstrating that actively cycling populations are confined predominantly to specific subsets.

### Multi-layered CD8^+^ T cell exhaustion and identification of a novel IEG^+^ effector memory state

To dissect functional states and differentiation relationships among CD8⁺ subsets, we examined marker expression and trajectory dynamics. The exhaustion transcription factors TOX, NR4A3 and BATF were co-enriched in both TOX⁺CD8_T and CD8_T_exhausted populations, confirming two transcriptionally distinct exhaustion-associated states within the same tumor (Fig. 2a). GZMK preferentially marked transitional and effector memory populations, GZMA showed intermediate distribution, and GZMM exhibited the broadest expression across multiple states, indicating a graded rather than binary pattern of cytotoxic capacity (Fig. 2b). A discrete IEG⁺CD8_Tem population was identified by high expression of FOS, FOSB and JUN (Fig. 2c), immediate early gene transcription factors that are rapidly induced upon TCR engagement and serve as master regulators of T cell activation; a distinct population defined by this signature has not been reported previously in the human tumor microenvironment.

**Fig. 2.**
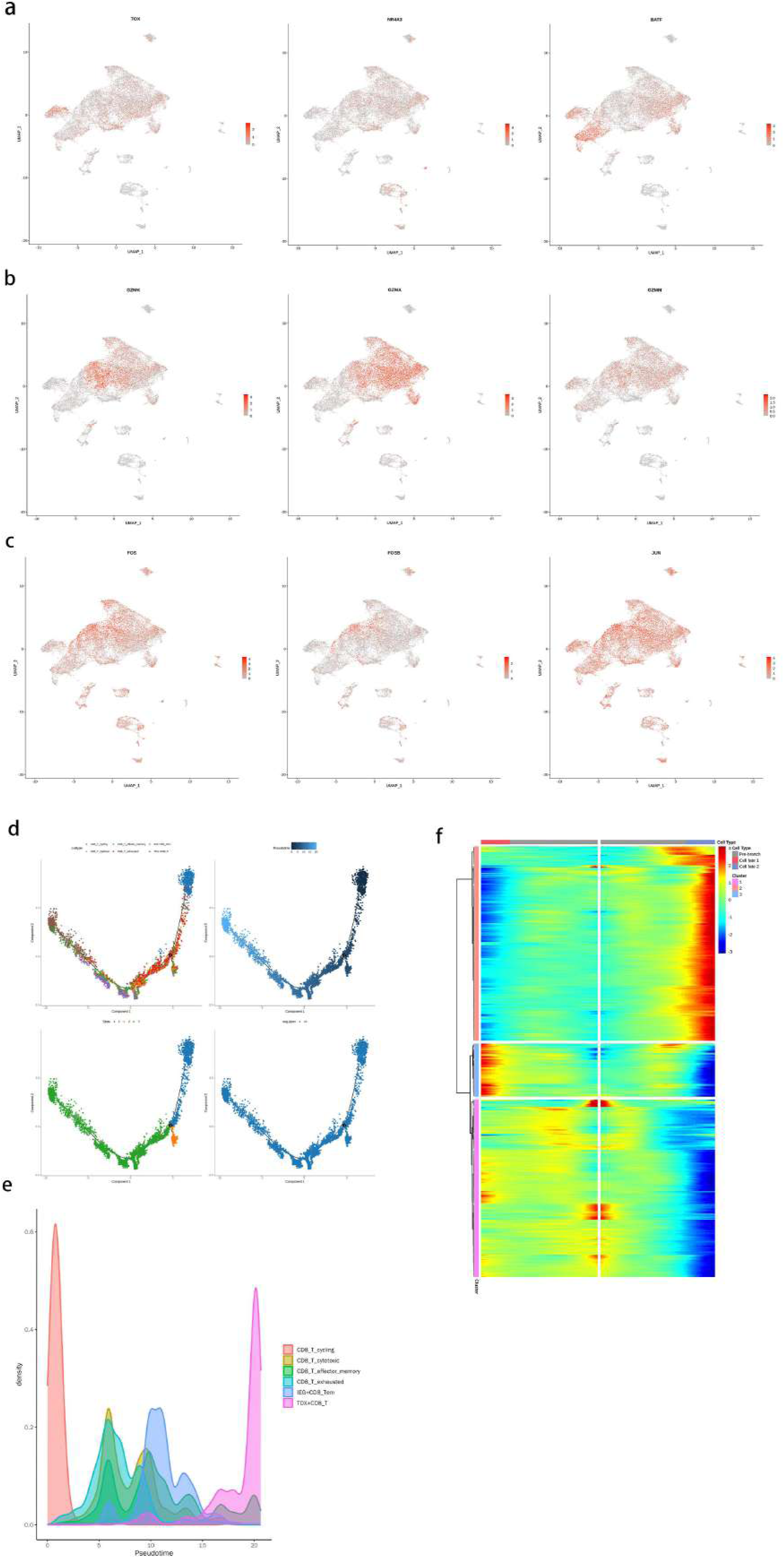
CD8⁺ T cell functional heterogeneity and differentiation trajectory. **a** Feature plots of exhaustion-associated transcription factors TOX, NR4A3 and BATF. **b** Feature plots of cytotoxic effector molecules GZMK, GZMA and GZMM. **c** Feature plots of immediate early gene markers FOS, FOSB and JUN, defining the IEG⁺CD8_Tem population. **d** Monocle2 pseudotime trajectory of six CD8⁺ T cell subsets, displayed as four panels colored by cell type (upper left), pseudotime (upper right), state (lower left) and sample identity (lower right). **e** Pseudotime density plot showing the temporal ordering of six CD8⁺ T cell states. **f** Branch point heatmap displaying gene expression dynamics across the exhaustion-versus-renewal fate decision.

Monocle2 pseudotime trajectory analysis revealed a branched differentiation architecture (Fig. 2d). Density analysis ordered the six states along this trajectory: CD8_T_cycling at the earliest position (∼1), CD8_T_effector_memory and CD8_T_exhausted at intermediate positions (∼5–7), CD8_T_cytotoxic at a later position (∼7–8), IEG⁺CD8_Tem at a more advanced position (∼10), and TOX⁺CD8_T at the terminal position (∼20) (Fig. 2e). Two aspects of this ordering are noteworthy. First, TOX⁺CD8_T cells occupied the terminal differentiation endpoint rather than canonically defined CD8_T_exhausted, indicating that exhaustion in HPV⁺ OPSCC extends beyond standard classifications. Second, IEG⁺CD8_Tem occupied a critical juncture between cytotoxic function and terminal exhaustion, suggesting that these cells represent recently antigen-restimulated T cells at a fate-decision point where the balance between sustained effector function and irreversible exhaustion is determined. Branch point analysis identified three gene clusters governing this exhaustion-versus-renewal decision (Fig. 2f), providing candidate molecular targets for redirecting T cell differentiation away from terminal dysfunction.

### Active Treg differentiation with comprehensive immunosuppressive receptor expression

Treg identity was validated by co-expression of FOXP3, IL2RA (CD25), CTLA4, TNFRSF4 (OX40), TNFRSF18 (GITR) and BATF, all colocalized to Treg and Treg_Cycling regions on UMAP (Fig. 3a). The simultaneous expression of co-inhibitory and co-stimulatory receptors on these Tregs identifies multiple surface molecules amenable to combinatorial antibody-based targeting. Monocle2 pseudotime analysis placed CD4_T_exhausted at early pseudotime (∼1) and Treg_Cycling at late pseudotime (∼12–13) (Fig. 3b–c), indicating active intratumoral Treg differentiation rather than passive peripheral recruitment. Monocle3 independently confirmed Treg as a terminal endpoint (Fig. 3d), and branch heatmaps revealed fate-decision gene modules governing the divergence between exhaustion and regulatory commitment (Fig. 3e). The shared expression of BATF across both Tregs and exhausted CD8⁺ T cells points to overlapping transcriptional circuits underlying immunosuppression in the CD4⁺ and CD8⁺ compartments.

**Fig. 3.**
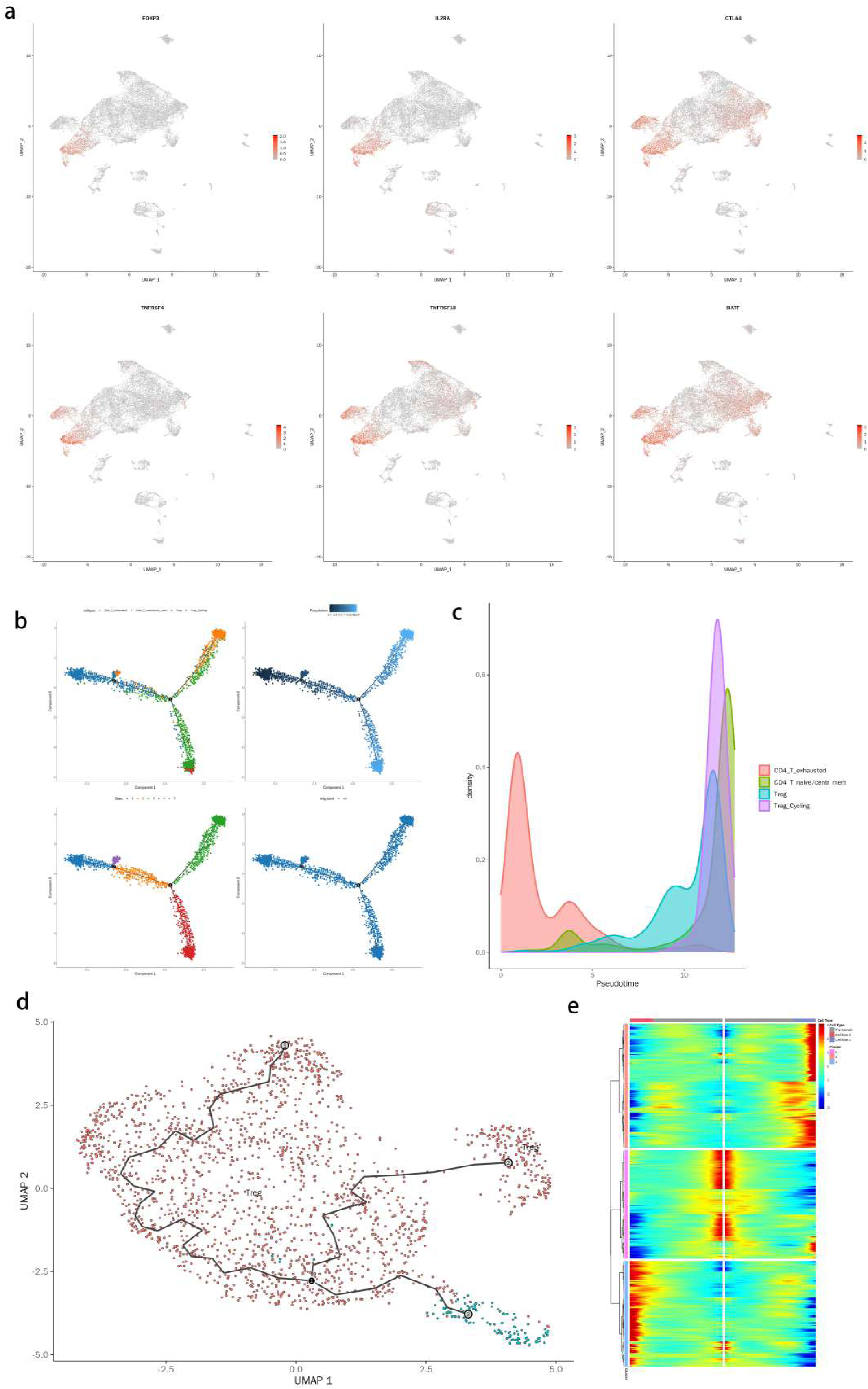
CD4⁺ T cell compartment and Treg differentiation. **a** Feature plots of Treg markers FOXP3, IL2RA (CD25), CTLA4, TNFRSF4 (OX40), TNFRSF18 (GITR) and BATF. **b** Monocle2 pseudotime trajectory of four CD4⁺ T cell subsets. **c** Pseudotime density plot. **d** Monocle3 trajectory analysis confirming Treg as a terminal endpoint. e Branch point heatmap of CD4⁺ fate-decision genes.

### Immunometabolic compartmentalization and intercellular communication

Feature plot analysis revealed compartmentalized metabolic gene expression (Fig. 4a–c). LDHA and SLC16A3 (MCT4), responsible for lactate production and export, were enriched in cancer cells, whereas SLC16A1 (MCT1) and LDHB, mediating lactate uptake and reverse conversion to pyruvate, were preferentially expressed in T cells. This complementary pattern supports directional lactate flux from glycolytic cancer cells to T cells, where imported lactate may reinforce exhaustion. KMT2D, CXCL14, TGFB2, VCAN, MMP10 and ACKR3 showed distinct compartmentalization across cancer, immune and stromal populations, with dot plot analysis confirming these patterns across all 24 cell types (Fig. 4d).

**Fig. 4.**
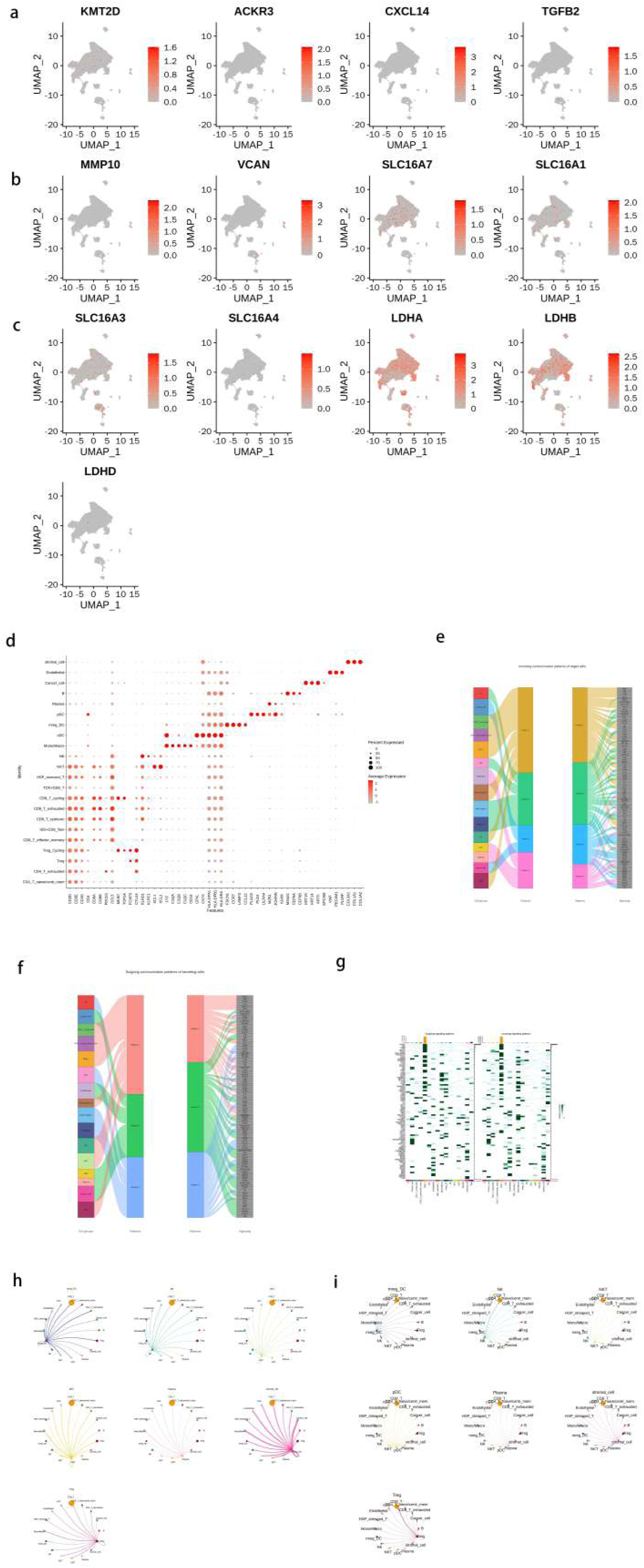
Immunometabolic landscape and intercellular communication. **a** Feature plots of microenvironmental signaling molecules: KMT2D, ACKR3, CXCL14 and TGFB2. **b** Feature plots of extracellular matrix and metabolic genes: MMP10, VCAN, SLC16A7 (MCT7) and SLC16A1 (MCT1). **c** Feature plots of glycolytic and lactate metabolism genes demonstrating compartmentalized expression. **d** Dot plot of key gene expression across all 24 cell types. **e** CellChat incoming communication patterns. **f** CellChat outgoing communication patterns. **g** Signaling role heatmap. **h** Individual signaling pathway circle plots. **i** Additional pathway interaction networks.

CellChat identified four outgoing and four incoming signaling patterns (Fig. 4e–f). Pattern 1, dominated by cancer cells and CD8⁺ T cells, mediated MHC-I/MIF/CXCL signaling reflecting antigen recognition; Pattern 3, driven by NK cells, pDC and mreg_DC, engaged PD-L1/BTLA/LIGHT checkpoint pathways. Cancer cells dominated outgoing MHC-I and MIF signals, while Tregs were principal recipients of ICOS/IL2/CD40 signals (Fig. 4g). mreg_DC served as a central signaling hub bridging recognition and suppression (Fig. 4h–i). The concurrent wiring of activation and inhibitory signals within the same network explains why single-agent ICB proves insufficient: blocking one checkpoint axis leaves parallel suppressive pathways intact.

### Spatial transcriptomics confirms tumor architecture and peripheral immune exclusion

Visium spatial transcriptomics with RCTD deconvolution mapped cell type distributions across an adjacent tissue section (Fig. 5a). Cancer cells dominated the spatial landscape (∼24,500/25,000 spots; Fig. 5b), with immune populations forming focal niches confined to the tumor periphery, consistent with an immune-excluded phenotype. Sub-clustering of non-cancer spots (n = 487) by t-SNE (Fig. 5c) and UMAP (Fig. 5d) resolved seven populations: macrophages (n = 142), monocytes (n = 89), dendritic cells (n = 67), CD8⁺ T cells (n = 58), CD4⁺ T cells (n = 52), neutrophils (n = 45) and NK cells (n = 34). Given the limited spot numbers, this sub-clustering serves as exploratory spatial annotation. KLF7 exhibited focal spatial expression consistent with a localized neural-associated microenvironment (Fig. 5e), and cancer cell deconvolution scores revealed heterogeneous tumor density with reduced cancer cell proportion corresponding to immune-infiltrated areas (Fig. 5f). The spatial juxtaposition of peripheral immune niches with metabolically active cancer cores thus explains how dense global infiltration coexists with functional immune exclusion at the local level.

**Fig. 5.**
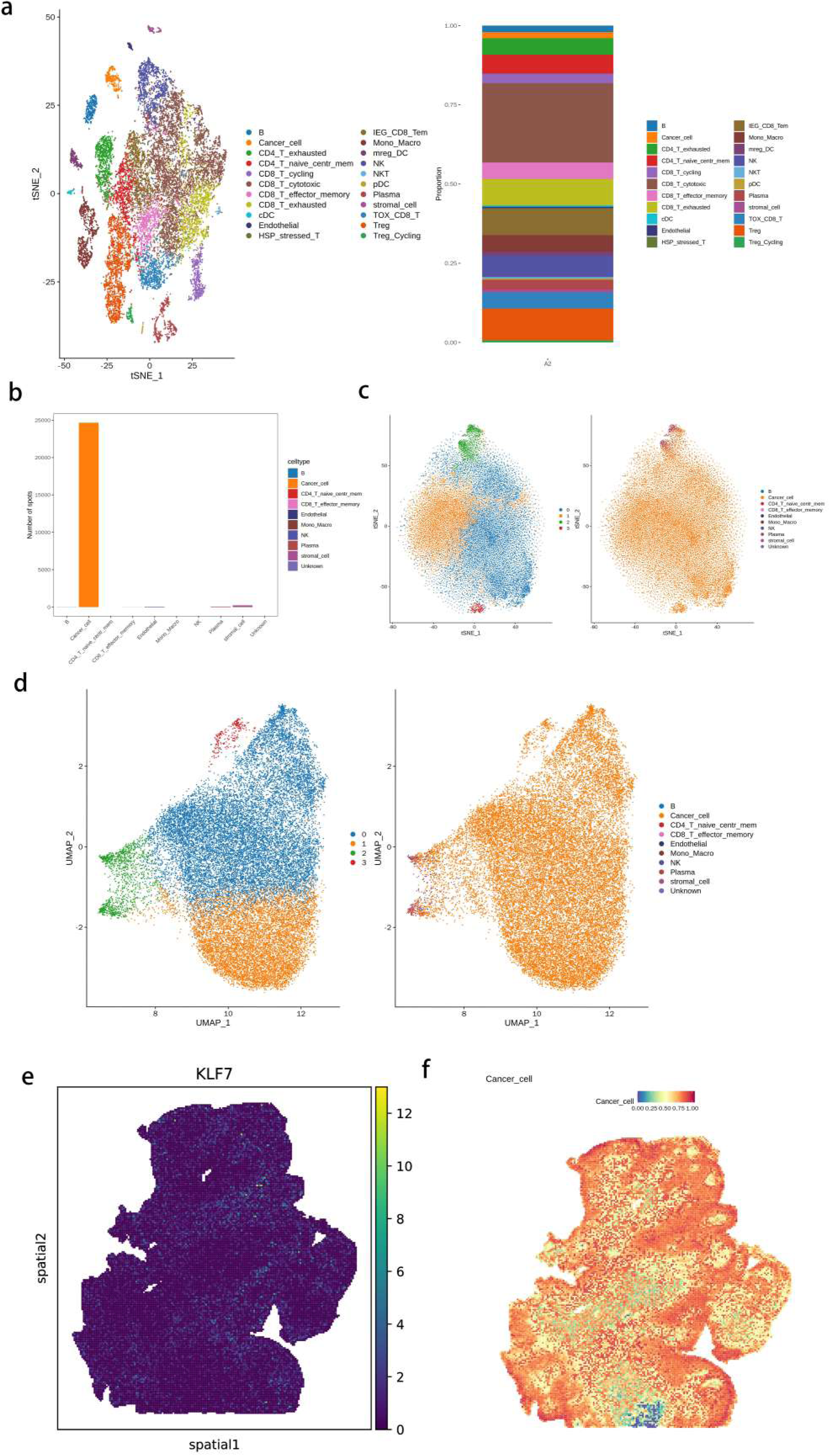
Spatial transcriptomics and cell type deconvolution. **a** Left: t-SNE of Visium spatial spots. Right: stacked bar plot of cell type proportions. **b** Bar plot of spot counts per primary cell type. **c** Sub-clustering of non-cancer spots by t-SNE. **d** Sub-clustering by UMAP. **e** Spatial expression map of KLF7. **f** Cancer cell deconvolution score mapped spatially.

### KMT2D-expressing cancer cells upregulate DNA repair and Wnt signaling

Eighteen genes were significantly upregulated in KMT2D-positive cancer cells. KEGG enrichment revealed concentration in base excision repair (PARP2; fold enrichment = 22.77), complement cascades (C1S) and transcription machinery (CDK19) (Fig. 6d). GO analysis demonstrated enrichment in β-catenin–TCF complex assembly (KMT2D), heterochromatin formation (KMT2D, BAHD1) and DNA ADP-ribosylation (PARP2) (Fig. 6e). Gene-function networks linked KMT2D, PARP2, C1S, CDK19 and BAHD1 to chromatin silencing complex and mitochondrial degradosome compartments (Fig. 6f–g). The coordinated upregulation of PARP2 and Wnt/β-catenin components links epigenetic remodeling to DNA repair capacity and stemness maintenance, suggesting selective vulnerability to PARP inhibitor therapy in KMT2D-expressing tumors.

**Fig. 6.**
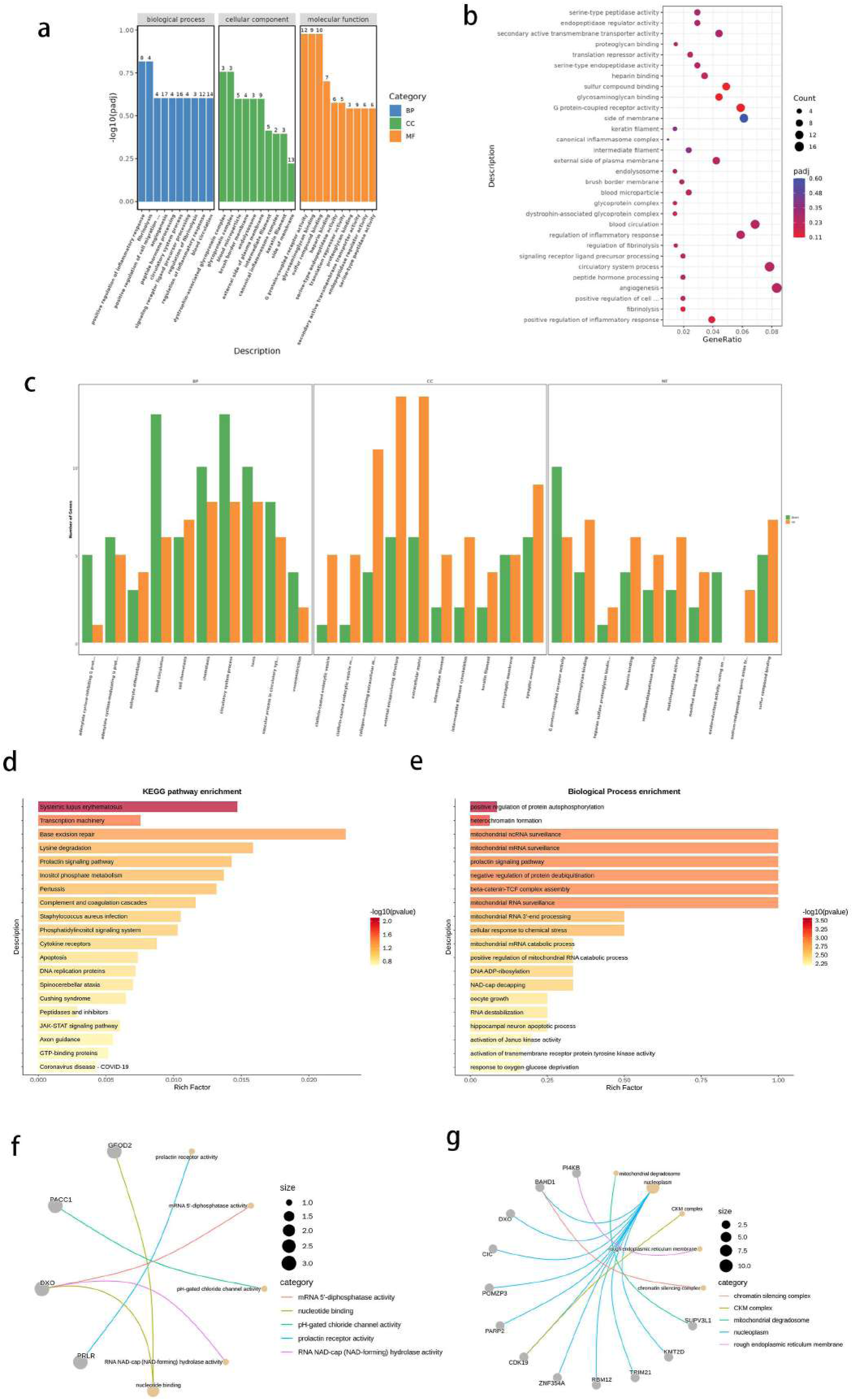
KMT2D functional characterization: CRISPR-Cas9 validation and single-cell enrichment analysis. **a** GO enrichment for CRISPR-Cas9 KMT2D-knockout comparison. **b** GO enrichment dot plot. **c** GO enrichment for an additional KMT2D-knockout comparison. **d** KEGG pathway enrichment of differentially expressed genes between KMT2D-expressing and KMT2D-non-expressing cancer cells. **e** GO Biological Process enrichment from single-cell KMT2D analysis. **f** Gene-function network (cnet) plot for Molecular Function terms. **g** Gene-function network (cnet) plot for Cellular Component terms.

### CRISPR-Cas9 knockout of KMT2D validates immune-regulatory functions and identifies KLF7 as a downstream target

Two independent KMT2D-knockout clones (KO1, KO2) were generated in SCC090 and characterized by RNA-seq. GO enrichment of the first knockout comparison revealed vascular process regulation, chemotaxis and extracellular matrix remodeling, with both upregulated and downregulated genes contributing, indicating bidirectional transcriptional reprogramming upon KMT2D loss (Fig. 6a). The second comparison showed enrichment in inflammasome complex assembly, inflammatory response and angiogenesis (Fig. 6b–c). The inter-clone comparison revealed enrichment specifically in lymphocyte and T cell proliferation regulation, demonstrating that graded KMT2D disruption differentially impacts immune-regulatory programs and providing functional evidence that tumor-intrinsic KMT2D modulates cancer cell capacity to regulate adaptive immune cell proliferation.

Western blot in both SCC090 and SCC154 showed that KMT2D knockout led to concurrent reduction of KMT2D, KLF7 and H3K4me1 protein levels (Fig. 7g). The concurrent loss of KLF7 and H3K4me1 establishes KMT2D as an upstream regulator of KLF7 through enhancer-mediated H3K4 monomethylation. Reproducibility across two independent HPV⁺ cell lines strengthens this regulatory relationship and bridges the epigenetic and neural microenvironment axes identified in this study.

**Fig. 7.**
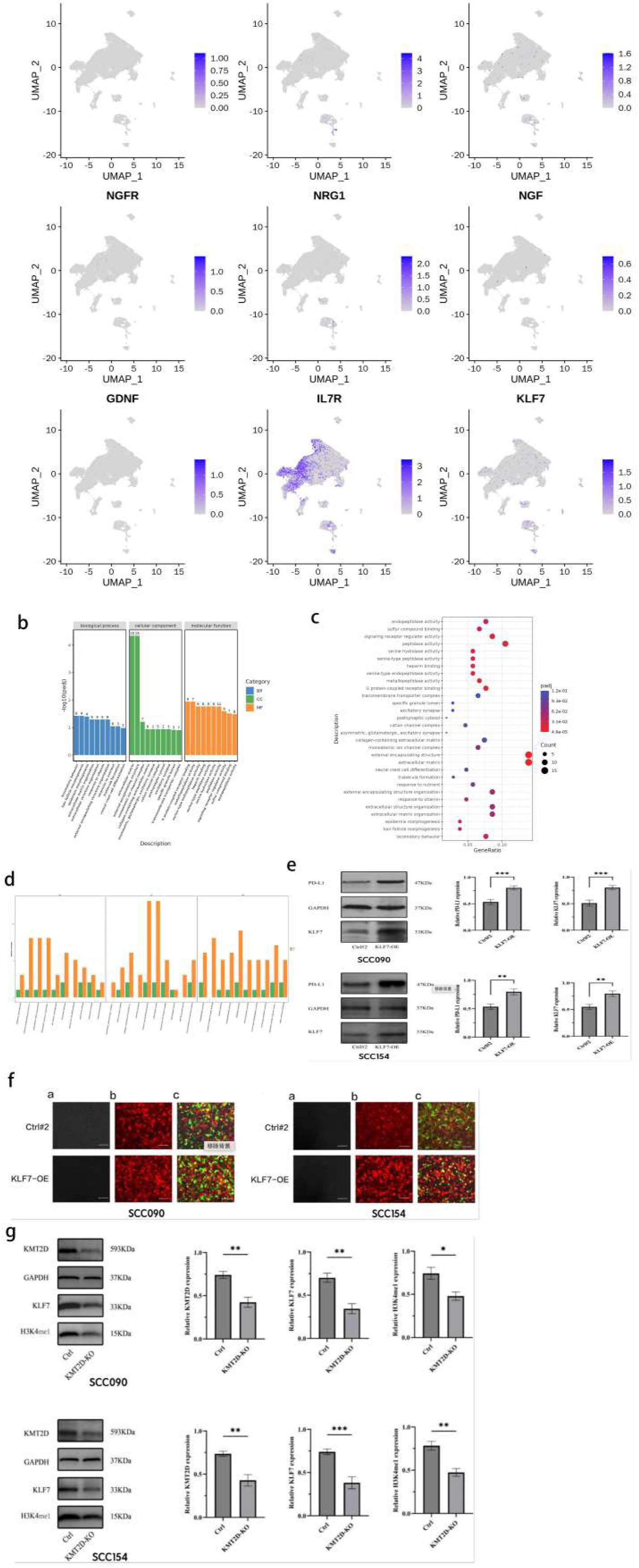
KLF7 functional characterization: neural differentiation, PD-L1 regulation, and KMT2D-KLF7 regulatory axis. **a** Feature plots of neural lineage markers. **b** GO enrichment for KLF7 overexpression versus control. **c** GO enrichment dot plot for KLF7 overexpression. **d** Directional enrichment bar plot. **e** Western blot of PD-L1 and KLF7 in control and KLF7-overexpressing cells. P < 0.01; P < 0.001. **f** Immunofluorescence of control and KLF7-overexpressing cells. **g** Western blot of KMT2D, KLF7 and H3K4me1 in control and KMT2D-knockout cells. P < 0.05; **P < 0.01.

### KLF7 overexpression drives neural crest differentiation, ECM remodeling, and PD-L1 upregulation

scRNA-seq identified a distinct population expressing neural markers S100B, NGFR, NRG1, NGF, GDNF and TUBB3 (Fig. 7a). KLF7 overexpression in both SCC090 and SCC154 induced GO enrichment in extracellular matrix organization (Padj < 10⁻⁴), neural crest cell differentiation, epidermis morphogenesis and locomotory behavior, with cellular component and molecular function analyses confirming ECM dominance and metallopeptidase activity upregulation (Fig. 7b–d). The neural crest enrichment validates the biological identity of the S100B⁺/NGFR⁺ population and establishes KLF7 as a driver of neural and Schwann-like cell programs within HPV⁺ OPSCC. Western blot revealed substantial PD-L1 upregulation upon KLF7 overexpression in both cell lines (Fig. 7e), confirmed by immunofluorescence (Fig. 7f). Together with the KMT2D–KLF7 regulatory relationship (Fig. 7g), these data define a KMT2D–KLF7–PD-L1 axis linking epigenetic remodeling to both neural crest-like reprogramming and immune checkpoint expression. These transcriptomic findings suggest KLF7 may link neural-associated populations to perineural invasion, although direct functional validation through dorsal root ganglion co-culture or in vivo assays will be required.

### Multiplex immunofluorescence validates key findings across independent HPV^+^ OPSCC cases

Five-marker mIF (DAPI/CD3/KMT2D/FOXP3/CK19) on independent HPV⁺ OPSCC cases confirmed four consistent patterns at the protein level (Fig. 8). Low-magnification imaging revealed CK19⁺ cancer cell nests surrounded by CD3⁺ T cell-rich stromal zones (Fig. 8a), while high-magnification views demonstrated scattered FOXP3⁺ Tregs and heterogeneous KMT2D expression in cancer nests (Fig. 8b). CD3⁺ T cell density reached 7,616.6 cells/mm² (Fig. 8c–d), confirming the T cell-dominant microenvironment. FOXP3⁺ Tregs accounted for 67.6% of CD3⁺ T cells (Fig. 8e), and 53.6% of CK19⁺ cancer cells co-expressed KMT2D (Fig. 8f). Peripheral immune niche distribution was consistent across specimens, recapitulating the spatial exclusion phenotype. These protein-level data across independent patients confirm that the immune heterogeneity, Treg distribution, KMT2D expression patterns and spatial organization are reproducible features of HPV⁺ OPSCC.

**Fig. 8.**
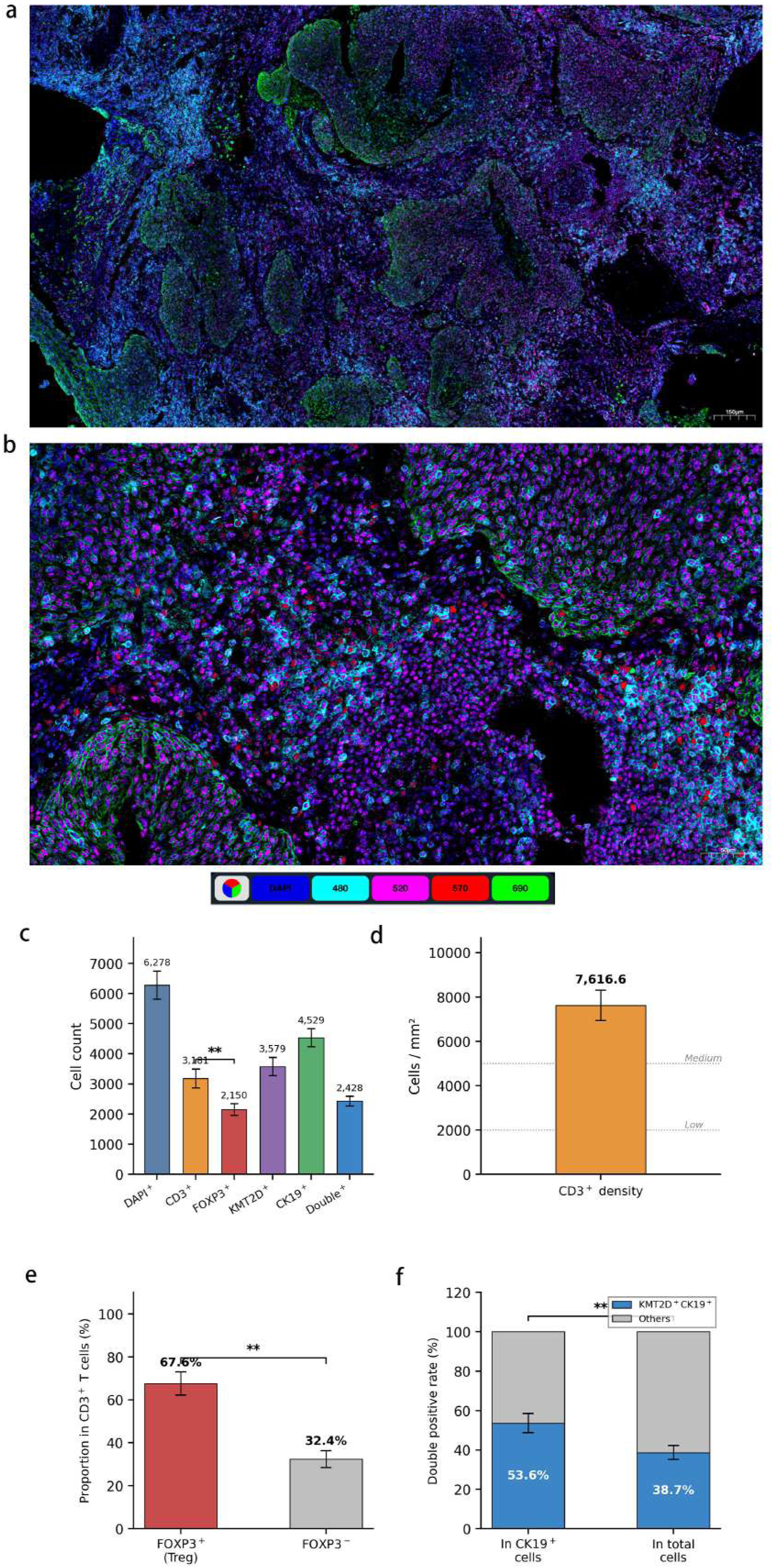
Multiplex immunofluorescence validation across independent HPV^+^ OPSCC cases. **a** Low-magnification mIF image. Scale bar: 150 μm. **b** High-magnification view of the tumor–immune interface. **c** Quantification of cell counts. **d** CD3⁺ T cell density. **e** Proportion of FOXP3⁺ versus FOXP3⁻ cells among CD3⁺ T cells. **f** KMT2D⁺CK19⁺ double-positive rate.

## Discussion

The tumor immune microenvironment of HPV-positive OPSCC presents a paradox with direct clinical consequences: despite one of the densest lymphocytic infiltrates among solid tumors, fewer than 20% of patients achieve durable responses to immune checkpoint blockade.^8,9^ Resolving this paradox requires understanding of immune cell heterogeneity, spatial organization, metabolic context and tumor-intrinsic molecular programs at single-cell resolution. Here, we provide an integrated single-cell, spatial and functionally validated atlas of the HPV⁺ OPSCC microenvironment, showing that ICB resistance arises from the convergence of multi-layered T cell exhaustion, immunometabolic compartmentalization, redundant intercellular signaling and a tumor-intrinsic KMT2D-KLF7-PD-L1 regulatory axis. Although the single-cell and spatial analyses derive from a single discovery specimen, multi-case protein-level validation, functional experiments in two independent cell lines and the biological coherence across orthogonal analytical approaches provide a framework connecting molecular mechanisms to actionable therapeutic strategies.

The precise temporal ordering of six CD8⁺ T cell functional states along a branched pseudotime trajectory represents a central finding. The prevailing model of T cell exhaustion posits relatively linear progression from progenitor-exhausted to terminally exhausted states defined by PD-1, TIM-3, LAG-3 and TOX co-expression.^11–14^ Our data challenge this framework in two ways. First, TOX⁺CD8_T cells, rather than canonically defined CD8_T_exhausted, occupied the terminal trajectory position, indicating that exhaustion in HPV⁺ OPSCC proceeds through an extended differentiation program beyond standard classifications. This carries practical implications for biomarker development, as strategies aimed at reinvigorating exhausted T cells must account for the possibility that the true terminal state may be missed by conventional marker panels. Second, the identification of a discrete IEG⁺CD8_Tem population at an intermediate position between cytotoxic function and terminal exhaustion represents a fate-decision checkpoint not previously characterized in the human tumor microenvironment. The high expression of FOS, FOSB and JUN in this population indicates recent TCR engagement and active transcriptional remodeling. Given that c-Jun overexpression confers exhaustion resistance in CAR-T cells,^16^ the IEG⁺CD8_Tem population may represent the cellular compartment most amenable to therapeutic rescue through checkpoint blockade or adoptive cell therapy. Branch point analysis further identified three gene clusters governing the exhaustion-versus-renewal fate decision, offering candidate targets for interventions designed to tip the balance toward sustained effector function. Single-cell TCR sequencing and ex vivo restimulation assays will be required to determine whether IEG⁺CD8_Tem cells retain clonal expansion capacity upon checkpoint release.

If the CD8⁺ trajectory defines the effector arm of the immune response, the Treg compartment defines its suppressive counterpart. Tregs co-expressed FOXP3, CTLA4, OX40 and GITR with active intratumoral cycling, and pseudotime analysis demonstrated intratumoral differentiation rather than passive peripheral recruitment. This is consistent with reports that PD-1 blockade can paradoxically amplify Treg numbers and suppressive activity in HNSCC,^17^ suggesting that single-agent anti-PD-1 therapy may inadvertently strengthen the immunosuppressive programs it aims to overcome. The co-expression of co-inhibitory (CTLA-4) and co-stimulatory (OX40, GITR) receptors provides a rationale for combining anti-CTLA-4 with agonistic anti-OX40 to deplete or reprogram Tregs while unleashing CD8⁺ effector function. mIF confirmed that FOXP3⁺ cells constituted 67.6% of the CD3⁺ T cell infiltrate (Fig. 8e), a proportion that exceeds the 5–30% range typically reported for HNSCC and warrants careful interpretation. Several factors may contribute: the mIF quantification was derived from representative high-magnification fields at the tumor–stroma interface where Treg density is highest, rather than whole-slide analysis; the FOXP3 positivity threshold in TSA-based mIF systems yields higher sensitivity than conventional immunohistochemistry; and activated human CD4⁺ T cells can transiently upregulate FOXP3 without acquiring suppressive function, which our panel lacking CD25/CD127 could not distinguish. Nonetheless, scRNA-seq data independently confirmed a substantial Treg and Treg_Cycling population with co-expression of the full immunosuppressive marker panel, supporting the biological significance of the Treg compartment regardless of the precise mIF-derived proportion. Future studies incorporating CD25⁺CD127⁻ co-staining will be needed to refine Treg quantification.

What prevents these abundant but dysfunctional T cells from accessing and eliminating cancer cells? The immunometabolic compartmentalization identified here offers one explanation. Complementary expression of LDHA and SLC16A3 (MCT4) in cancer cells versus LDHB and SLC16A1 (MCT1) in T cells defines a directional lactate flux model in which glycolytic cancer cells export lactate, which is then imported by neighboring T cells. Lactate uptake through MCT11 enforces T cell dysfunction by disrupting NAD⁺/NADH redox balance,^21^ while lactate-driven MOESIN lactylation enhances TGF-β signaling in Tregs.^22^ Our spatial transcriptomic data support this model at the tissue level: immune cell niches were confined to the tumor periphery, spatially separated from the metabolically active cancer cell core, consistent with a lactate-mediated biochemical barrier that restricts effective immune cell penetration into the tumor parenchyma. This spatial immune exclusion persists despite the overall T cell-rich character of HPV⁺ OPSCC, explaining at the tissue level why global measures of immune infiltration fail to predict ICB response. Therapeutic disruption of this metabolic barrier through LDHA or MCT4 inhibitors, combined with checkpoint blockade, represents a strategy that directly addresses the spatial disconnect between immune cell presence and function.

CellChat analysis reveals an additional layer of resistance at the intercellular signaling level. Four distinct communication patterns demonstrated that immune activation (MHC-I, MIF, CXCL signaling in Pattern 1) and immune suppression (PD-L1, BTLA, LIGHT engagement in Pattern 3) coexist within the same signaling network. Cancer cells simultaneously present antigens through MHC-I and engage checkpoint pathways, while Tregs receive co-stimulatory signals (ICOS, IL2, CD40) that maintain their suppressive function. mreg_DC emerged as a central signaling hub bridging recognition and suppression, a dual role consistent with the regulatory function attributed to LAMP3⁺ mature dendritic cells in other tumor types. This concurrent wiring of activation and inhibition explains mechanistically why single-agent ICB is insufficient: blocking one checkpoint axis leaves parallel suppressive pathways intact. Effective immunotherapy will likely require simultaneous disruption of multiple pathways-for instance, combining anti-PD-1 with anti-TIGIT or anti-BTLA-to dismantle the redundant inhibitory architecture.

The KMT2D-KLF7-PD-L1 axis integrates what initially appeared to be independent observations into a unified tumor-intrinsic regulatory framework. At the single-cell level, KMT2D-expressing cancer cells exhibited coordinated upregulation of PARP2 (base excision repair; fold enrichment = 22.77), C1S (complement cascade) and β-catenin-TCF complex assembly genes, linking epigenetic remodeling to both DNA repair capacity and Wnt-driven stemness maintenance. CRISPR-Cas9 knockout experiments moved these observations from correlation to causation. The three-way comparison across wild-type, KO1 and KO2 cells revealed a hierarchical functional architecture: one comparison demonstrated vascular remodeling and inflammasome activation, while the inter-clone comparison uncovered regulation of lymphocyte and T cell proliferation, indicating that KMT2D functions as a dose-dependent rheostat for immune-regulatory gene programs. Concurrent reduction of KLF7 protein and H3K4me1 levels upon KMT2D depletion in both SCC090 and SCC154 (Fig. 7g) demonstrates that KMT2D maintains KLF7 expression through enhancer-mediated epigenetic regulation, placing KMT2D at the apex of a regulatory hierarchy extending from chromatin remodeling through neural microenvironment programming to immune checkpoint control.

KLF7 overexpression drove neural crest cell differentiation and ECM remodeling while simultaneously upregulating PD-L1 protein in both cell lines (Fig. 7e–f), revealing a dual function that connects neural programming directly to immune evasion. The induction of neural crest differentiation validates the biological identity of the S100B⁺/NGFR⁺ population and establishes KLF7 as a functional driver of Schwann-like cell programs.^38,40^ Concurrent ECM remodeling and metallopeptidase activation define a transcriptional program consistent with tissue remodeling associated with perineural spread, although direct functional evidence from dorsal root ganglion co-culture or in vivo perineural invasion (PNI) assays is lacking; these PNI-related findings should be considered hypothesis-generating. To explore whether KLF7 directly regulates PD-L1 transcription, we scanned the 2 kb region upstream of the CD274 transcription start site using the JASPAR database and identified two putative KLF-family binding motifs (CACCC-box elements) at positions-847 and-1,234 (matrix similarity scores 0.87 and 0.82). Whether KLF7 physically occupies these sites remains to be determined by ChIP-qPCR or luciferase reporter assays; alternatively, PD-L1 upregulation may occur indirectly through KLF7-mediated activation of IFN-γ response genes or JAK–STAT signaling. From a therapeutic perspective, the convergence of PARP2 upregulation, KLF7-dependent PD-L1 modulation and neural crest reprogramming within a single regulatory cascade provides a multi-pronged rationale for targeting KMT2D-associated vulnerabilities: PARP inhibition would exploit the DNA repair dependency of KMT2D-expressing cancer cells while simultaneously disrupting the epigenetic maintenance of the KLF7-PD-L1 immune evasion program. The hierarchical organization of this axis is summarized in Fig. 9.

**Fig. 9.**
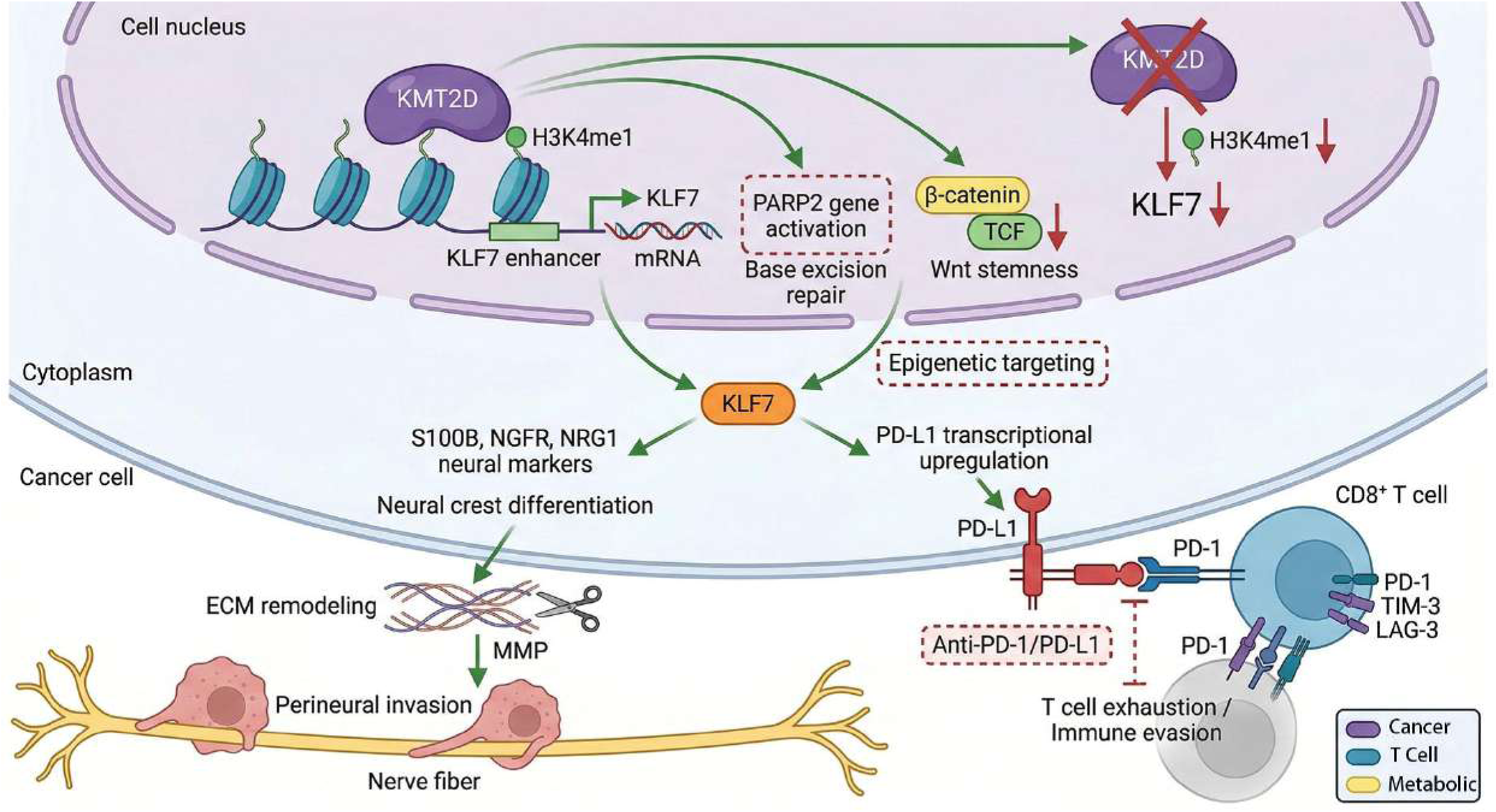
Schematic illustration of the KMT2D-KLF7-PD-L1 regulatory axis in HPV-positive oropharyngeal squamous cell carcinoma. KMT2D catalyzes H3K4me1 at enhancer elements, activating KLF7 transcription as well as PARP2 and β-catenin–TCF complex assembly genes. KLF7 drives two functionally distinct downstream programs: neural crest differentiation with ECM remodeling; and PD-L1 upregulation enabling immune evasion. Red dashed boxes indicate therapeutic intervention points.

mIF validation across independent cases strengthens the generalizability of our findings. Beyond qualitative patterns (Fig. 8a–b), quantitative analysis provided specific metrics: CD3⁺ T cell density of 7,616.6 cells/mm² confirmed the T cell-dominant microenvironment; KMT2D⁺CK19⁺ double-positive rate of 53.6% among cancer cells established the prevalence of KMT2D expression in the epithelial compartment; and peripheral immune niche distribution was consistent across specimens. Protein-level validation using an orthogonal technique in an independent cohort addresses a common limitation of single-specimen single-cell studies and supports the biological relevance of the cell states, spatial organization and molecular programs described here.

Several limitations should be acknowledged. The scRNA-seq and spatial analyses derived from a single discovery specimen, constraining assessment of inter-patient heterogeneity in cell type proportions, trajectory dynamics and spatial organization. The quantitative features reported should be interpreted as representative of the specimen analyzed rather than as population-level estimates. Multi-patient scRNA-seq studies with matched spatial profiling and clinical outcome data are needed to determine the prevalence and clinical significance of the cell states identified. Sub-clustering of ∼500 non-cancer Visium spots into seven populations yielded small cluster sizes (34–142 spots per cluster), limiting statistical power; this spatial annotation should be considered exploratory, and higher-resolution methods such as MERFISH or CosMx would enable more robust spatial interaction analysis. The PNI conclusions rest on transcriptomic enrichment without direct functional assays and should be considered hypothesis-generating. The mechanistic basis for KLF7-mediated PD-L1 regulation-whether through direct promoter binding or indirect pathways-awaits experimental clarification. Prospective clinical studies correlating our molecular features with ICB treatment outcomes are needed to translate these findings into biomarker-guided strategies.

In summary, the integrated atlas presented here reveals that ICB failure in HPV⁺ OPSCC arises from converging mechanisms operating across biological scales: a branched CD8⁺ T cell exhaustion trajectory with a fate-decision checkpoint at the IEG⁺ stage; active intratumoral Treg differentiation maintaining local immunosuppression; a metabolic barrier driven by compartmentalized lactate flux that spatially restricts immune cell function to the tumor periphery; redundant intercellular signaling that simultaneously engages immune recognition and checkpoint suppression; and a tumor-intrinsic KMT2D-KLF7-PD-L1 regulatory axis linking epigenetic remodeling to DNA repair vulnerability, neural microenvironment programming and immune checkpoint expression. On the basis of these findings, we propose a multi-target therapeutic framework: checkpoint combinations (anti-PD-1 with anti-TIGIT or anti-BTLA) to dismantle redundant suppressive signaling; Treg-directed agents (anti-CTLA-4 with agonistic anti-OX40) to interrupt immunosuppressive differentiation; metabolic interventions (LDHA or MCT4 inhibitors) to dissolve the lactate barrier and restore immune cell access to the tumor core; PARP inhibitors for KMT2D-expressing tumors to exploit DNA repair dependency while disrupting KLF7-driven programs; and investigation of the KMT2D-KLF7-PD-L1 axis as a candidate strategy to concurrently reduce perineural invasion risk and immune checkpoint-mediated evasion, pending functional validation of the PNI component. Prospective validation stratifying patients by KMT2D expression, KLF7-PD-L1 co-expression, IEG⁺CD8_Tem abundance, Treg cycling activity and spatial immune configuration will determine whether this biomarker-guided precision framework can improve outcomes for HPV⁺ OPSCC patients who currently derive limited benefit from immunotherapy.

## Methods

### Patient samples and ethics

The discovery cohort consisted of a surgically resected HPV⁺ OPSCC specimen collected at the Affiliated Hospital of Qingdao University for scRNA-seq and Visium spatial transcriptomics. The validation cohort comprised independent HPV⁺ OPSCC cases for multiplex immunofluorescence. HPV status was confirmed by p16 immunohistochemistry and HPV DNA detection. Written informed consent was obtained from each patient prior to tissue collection. This study was approved by the Ethics Committee of the Affiliated Hospital of Qingdao University and conducted in accordance with the 1964 Helsinki Declaration and its later amendments. Cell line experiments using SCC090 and SCC154 were performed in accordance with institutional biosafety regulations.

### Study design rationale

The discovery cohort was designed as a single-specimen deep-profiling study to maximize sequencing depth and enable characterization of rare cell populations that would be diluted in multi-patient pooled designs with shallower per-patient coverage. To address this inherent limitation, we implemented multi-layered validation: multiplex immunofluorescence across independent cases to confirm key cell populations and spatial organization at the protein level; and CRISPR-Cas9 knockout and overexpression experiments in two independent HPV⁺ cell lines to establish causal molecular relationships. This approach follows precedents established by landmark single-cell studies in the head and neck cancer field.^24^

### Single-cell RNA sequencing and data processing

Fresh tumor tissue was mechanically minced and enzymatically dissociated using collagenase IV and DNase I at 37 °C for 30 min with intermittent agitation. The resulting cell suspension was filtered through 70 μm and 40 μm cell strainers sequentially, and cell viability was assessed by trypan blue exclusion (>85% required). Single-cell capture and library preparation were performed using the 10x Genomics Chromium 3’ v3.1 system targeting ∼10,000 cells. Libraries were sequenced on an Illumina NovaSeq 6000 platform. Sequencing reads were aligned to the GRCh38-2024-A reference genome using Cell Ranger (10x Genomics).

Downstream analysis was performed using Seurat v4.x in R. Cells were filtered using the following criteria: minimum 500 and maximum 20,000 UMIs per cell; minimum 500 and maximum 8,000 genes detected per cell; maximum 20% mitochondrial gene content. Genes expressed in fewer than 3 cells were excluded. DoubletFinder was applied with an estimated doublet rate of 5% based on the 10x Genomics expected rate for the target cell recovery. After filtering, 23,119 cells were retained from an initial capture of ∼26,000 cells. SCTransform normalization was performed with regression of mitochondrial gene percentage. PCA was computed on the top 3,000 variable genes, and the first 30 principal components were selected based on elbow plot assessment. Louvain clustering was performed at a resolution of 0.8, and UMAP and t-SNE embeddings were computed with default Seurat parameters. Cell types were annotated based on canonical marker gene expression. Detailed marker gene lists are provided in Supplementary Methods.

### Pseudotime trajectory analysis

Monocle2 was applied separately to CD8⁺ T cells (6 subtypes) and CD4⁺ T cells (4 subtypes) using DDRTree dimensionality reduction to infer differentiation trajectories and identify branch points. Pseudotime density plots were generated to visualize the distribution of each subset along the trajectory. Monocle3 was used to independently validate CD4⁺ T cell trajectories and confirm terminal endpoints.

### Cell-cell communication analysis

CellChat v1.x was used to infer ligand–receptor interactions across all 24 cell types. Non-negative matrix factorization identified outgoing and incoming signaling patterns. Signaling role heatmaps and individual pathway interaction networks were computed to characterize the directionality and specificity of intercellular communication.

### Spatial transcriptomics

10x Visium spatial transcriptomics was performed on a formalin-fixed, paraffin-embedded section adjacent to the tissue used for scRNA-seq. Sequencing reads were processed using Space Ranger (10x Genomics). RCTD deconvolution was applied using the scRNA-seq atlas (22 cell types) as a reference to assign primary cell type identities to individual spatial spots. Non-cancer spots were extracted and sub-clustered independently to resolve immune and stromal populations within the spatial context.

### KMT2D differential expression analysis

Cancer cells from the scRNA-seq atlas were stratified into KMT2D-expressing and KMT2D-non-expressing groups. Differentially expressed genes were identified using the Wilcoxon rank-sum test with Benjamini–Hochberg FDR correction. GO (Biological Process, Cellular Component, Molecular Function), KEGG and Reactome pathway enrichment analyses were performed using clusterProfiler.

### Cell lines and culture conditions

Two HPV16-positive head and neck squamous cell carcinoma cell lines were used: SCC090 (UM-SCC-090, derived from a base of tongue squamous cell carcinoma) and SCC154 (UM-SCC-154, derived from a tonsillar squamous cell carcinoma). Cells were maintained in Dulbecco’s Modified Eagle’s Medium supplemented with 10% fetal bovine serum and 1% penicillin–streptomycin at 37 °C in a humidified atmosphere containing 5% CO₂. Both cell lines were used for CRISPR-Cas9 KMT2D knockout and KLF7 overexpression to ensure observed phenotypes were not cell line-specific. RNA-seq was performed on SCC090 for transcriptomic analysis, while Western blot and immunofluorescence validation were conducted in both cell lines.

### CRISPR-Cas9 KMT2D knockout and RNA-seq

KMT2D-knockout clones were generated in both SCC090 and SCC154 using the lentiviral CRISPR-Cas9 system. Single guide RNAs targeting distinct exons of KMT2D were designed and cloned into lentiCRISPR v2 (Addgene, #52961). Lentiviral particles were produced in HEK293T cells by co-transfection with psPAX2 (Addgene, #12260) and pMD2.G (Addgene, #12259) packaging plasmids using Lipofectamine 3000 (Thermo Fisher Scientific). Transduced cells were selected with puromycin (2 μg/mL) for 72 h, and single-cell clones were isolated by limiting dilution. Two independent knockout clones (KO1 and KO2) were confirmed by Western blot. Parental cells served as wild-type control. Detailed sgRNA sequences and knockout validation data are provided in Supplementary Methods.

For transcriptomic characterization, total RNA was extracted from SCC090 KO1, KO2 and wild-type cells using TRIzol reagent, and RNA integrity was verified by Agilent Bioanalyzer (RIN > 7.0). Libraries were prepared using the NEBNext Ultra II RNA Library Prep Kit for Illumina (New England Biolabs) and sequenced on an Illumina NovaSeq 6000 platform with paired-end 150 bp reads. Reads were aligned to the GRCh38 reference genome using HISAT2, and gene expression was quantified using featureCounts. Differential expression analysis was performed for three comparisons (KO1 vs wild-type, KO2 vs wild-type, KO2 vs KO1) using DESeq2 with FDR < 0.05 and |log₂ fold change| > 1 as significance thresholds. GO enrichment was performed using clusterProfiler with FDR correction.

### KLF7 overexpression and RNA-seq

The full-length human KLF7 coding sequence (NM_003709.4) was amplified by PCR and cloned into the pLVX-EF1α-IRES-Puro lentiviral expression vector (Clontech) between EcoRI and BamHI restriction sites. The construct was verified by Sanger sequencing. Lentiviral particles were produced in HEK293T cells as described above, and both SCC090 and SCC154 cells were transduced with KLF7-overexpression or corresponding empty vector control lentivirus. Successfully transduced cells were selected with puromycin (2 μg/mL) for 72 h. Overexpression efficiency was confirmed by quantitative real-time PCR and Western blot at 48 h post-selection. RNA-seq and differential expression analysis were performed on SCC090 cells as described above.

### Western blot

Total protein was extracted using RIPA lysis buffer supplemented with protease and phosphatase inhibitor cocktail. Equal amounts of protein were separated by SDS-PAGE and transferred to PVDF membranes. Membranes were blocked with 5% non-fat milk in TBST for 1 h, then incubated overnight at 4 °C with primary antibodies: anti-KMT2D (Sigma-Aldrich, HPA035977, 1:500), anti-KLF7 (Abcam, ab197690, 1:1000), anti-H3K4me1 (Abcam, ab8895, 1:2000), anti-PD-L1 (Cell Signaling Technology, #13684, 1:1000) and anti-GAPDH (Proteintech, 60004-1-Ig, 1:5000) as loading control. HRP-conjugated secondary antibodies were applied for 1 h at room temperature. Protein bands were detected using enhanced chemiluminescence substrate (Millipore) and visualized by the ChemiDoc Imaging System (Bio-Rad). Densitometric quantification was performed using ImageJ software (NIH), with relative protein expression normalized to GAPDH. All experiments were performed in at least three independent biological replicates.

### Immunofluorescence

KLF7-overexpressing and control cells from both SCC090 and SCC154 were seeded on glass coverslips in 24-well plates. At 48 h post-transduction, cells were fixed with 4% paraformaldehyde for 15 min, permeabilized with 0.1% Triton X-100 for 10 min and blocked with 5% BSA in PBS for 1 h. Cells were incubated with anti-PD-L1 primary antibody (Cell Signaling Technology, #13684, 1:200) overnight at 4 °C, followed by Alexa Fluor 594-conjugated goat anti-rabbit secondary antibody (Thermo Fisher Scientific, A-11012, 1:500) for 1 h at room temperature. Nuclei were counterstained with DAPI (1 μg/mL). Images were acquired using a fluorescence microscope with consistent exposure settings across all conditions.

### Multiplex immunofluorescence

Formalin-fixed, paraffin-embedded sections (4 μm) from the validation cohort were subjected to sequential multiplex immunofluorescence staining using the Opal 7-Color Manual IHC Kit (Akoya Biosciences) with tyramide signal amplification. Primary antibodies and corresponding Opal fluorophores were applied in the following order: CD3 (Dako, A0452, 1:200, Opal 480), KMT2D (Sigma-Aldrich, HPA035977, 1:100, Opal 520), FOXP3 (Abcam, ab20034, 1:100, Opal 570) and CK19 (Abcam, ab52625, 1:200, Opal 690). Between each staining round, antigen retrieval was repeated in citrate buffer (pH 6.0) to strip the previous antibody complex while retaining the covalently bound TSA signal. Nuclei were counterstained with DAPI. Whole-slide imaging was performed at 20× magnification using the Vectra Polaris Automated Quantitative Pathology Imaging System (Akoya Biosciences). Spectral unmixing, cell segmentation and phenotyping were performed using inForm Tissue Analysis Software (Akoya Biosciences) with machine learning-based tissue and cell classification algorithms trained on representative regions of interest. Positivity thresholds for each marker were determined by comparison with single-stained controls and pathologist review. Cell density was calculated as the number of positive cells per mm² of analyzed tissue area.

### Statistical analysis

For single-cell differential expression, the Wilcoxon rank-sum test was used with Benjamini–Hochberg FDR correction. Pathway enrichment analyses employed hypergeometric tests with FDR correction. For bulk RNA-seq, DESeq2 was applied with FDR < 0.05 and |log₂ fold change| > 1. CellChat statistical significance was assessed using permutation-based P values. For Western blot densitometry and multiplex immunofluorescence quantification, statistical comparisons between two groups were performed using two-tailed unpaired Student’s t-test. Data are presented as mean ± standard deviation from at least three independent experiments. Statistical significance was defined as P < 0.05, P < 0.01 and P < 0.001. All computational analyses were performed in R v4.x with visualization by ggplot2, Seurat, Monocle2, Monocle3, CellChat and custom scripts.

## Supporting information

Graphical Abstract

## Data availability

The scRNA-seq and spatial transcriptomics data generated in this study will be deposited in a publicly accessible repository upon publication. All other data supporting the findings of this study are available from the corresponding authors upon reasonable request.

## Acknowledgements

This work was supported by the National Natural Science Foundation of China (Grant No. 82203418) and the Postdoctoral Innovation Project of Shandong Province (Grant No. SDCX-ZG-2024-00031).

## Author contributions

**K.W.** and **N.D**.: Conceptualization, Methodology, Formal analysis, Data curation, Visualization, Writing-original draft. **Y.T**.: Conceptualization, Literature curation, Writing-original draft, Formal analysis. **X.Y**. and **M.Y.**: Investigation, Data curation, Resources. **L.G.** and **S.J**.: Software, Formal analysis (bioinformatics), Visualization. **W.S.**: Supervision, Investigation, Writing-review & editing. **J.D**. and **L.W.**: Conceptualization, Supervision, Funding acquisition, Project administration, Writing-review & editing.

---

**Supplementary Fig. S1 Myeloid compartment validation.** Feature plots of LYZ, CST3, C1QB (top row); C1QA, C1QC, FCER1G (middle row); CD14 and IL1B (bottom row), confirming monocyte and macrophage annotation. C1QA, C1QB, and C1QC are co-expressed in the macrophage cluster, while CD14 and LYZ mark the broader monocyte/macrophage population.

**Supplementary Fig. S2 Additional T cell markers.** Feature plots of ITK (left) and NFKB1 (right). ITK (IL-2 inducible T cell kinase) confirms T cell lineage identity across all T cell subsets. NFKB1 expression confirms active NF-κB signaling, with enrichment in specific immune populations.

**Supplementary Fig. S3 Validation of the IEG+CD8_Tem population in independent public scRNA-seq datasets.** CD8+ T cells from GSE139324 (Cillo et al., Immunity 2020; 26 patients: 8 HPV+, 18 HPV−) and GSE164690 (Kürten et al., Nat. Commun. 2021; 18 patients: 6 HPV+, 12 HPV−) were integrated using Harmony, yielding 3,637 high-quality cells (1,141 HPV+; 2,496 HPV−).

**a** Data integration quality control. Left: Feature scatter plots of QC metrics (nCount_RNA, nFeature_RNA, percent.mt, percent.rp) colored by HPV status (HPV−, red; HPV+, teal); Pearson r values shown. Overlapping distributions confirm equivalent data quality between groups. Right: UMAP after Harmony integration colored by HPV status; interdigitation of both groups confirms successful batch correction. **b** CD8+ T cell extraction workflow. Left: UMAP of all 40 clusters after Harmony integration. Middle: Seven CD8+ clusters (g0, g9, g12, g13, g17, g22, g29; CD8A mean > 3) selected based on CD8A/CD8B/CD3D co-expression. Right: UMAP of the 3,637 extracted CD8+ T cells in isolation. **c** CD8+ T cell sub-clustering and annotation. Left: UMAP colored by six annotated subtypes corresponding to those in the discovery cohort: CD8_T_cytotoxic (GZMK, GZMA), CD8_T_effector_memory, CD8_T_exhausted (PDCD1, HAVCR2, LAG3), IEG+CD8_Tem (FOS, FOSB, JUN), TOX+CD8_T (TOX, NR4A3), and CD8_T_cycling (MKI67, TOP2A). Right: Dot plot of marker gene expression across subtypes; dot size = percent expressing, color = average scaled expression. **d** UMAP feature plots of FOS, FOSB, and JUN across integrated CD8+ T cells. All three IEG transcription factors are co-enriched in a discrete subpopulation corresponding to the IEG+CD8_Tem cluster (panel c), recapitulating the signature identified in the HPV+ OPSCC discovery cohort. **e** IEG expression quantification. Left: Violin plots of FOS, FOSB, and JUN across the six CD8+ subtypes; IEG+CD8_Tem shows the highest expression of all three markers. Middle: Cells in the top 30% of combined FOS/JUN score (≥2.365; n = 1,091; orange) are spatially concentrated within the IEG+CD8_Tem cluster. Right: CD8_T_exhausted and TOX+CD8_T cells shown for topological comparison; their spatial separation from the IEG+ cluster is consistent with the pseudotime trajectory positioning. **f** CD8+ T cell subtype proportions by HPV status (integrated datasets). Stacked bar plots for HPV+ (n = 1,141 cells, 14 patients) versus HPV− (n = 2,496 cells, 30 patients). Combined FOS/JUN score was significantly higher in HPV− specimens (median 1.828 vs. 1.452; Wilcoxon p = 2.74×10−17; FOS p = 1.26×10−5, JUN p = 3.46×10−17), indicating that IEG+CD8_Tem is a conserved state across both HPV+ and HPV− head and neck tumor contexts. **g** Re-analysis of GSE164690 (Kürten et al.). Left: UMAP of re-clustered CD8+ T cells colored by cluster identity. Right: FOS expression on the same UMAP; a discrete IEG-high subpopulation is identifiable, recapitulating the IEG+CD8_Tem cluster observed in GSE139324 and the discovery cohort. **h** Violin plots of FOS, FOSB, and JUN across CD8+ subtypes in GSE164690. IEG+CD8_Tem exhibits markedly elevated expression of all three markers, consistent with GSE139324 results and validating subtype identity in an independent cohort. **i** UMAP feature plots of FOS, FOSB, and JUN in GSE164690. All three IEG markers are co-enriched in a spatially discrete subpopulation, confirming dataset-independent reproducibility of the IEG+CD8_Tem transcriptional signature. **j** CD8+ T cell subtype proportions by HPV status in GSE164690. Stacked bar plots for HPV+ (n = 919 cells, 6 patients) versus HPV− (n = 1,436 cells, 12 patients). IEG+CD8_Tem cells are detectable in both groups, with higher FOS/JUN intensity in HPV− specimens (JUN p = 3.46×10−17). **k** Statistical comparison of combined FOS/JUN scores between HPV+ and HPV− CD8+ T cells. Distribution of IEG score [mean(FOS, JUN)] for HPV+ (n = 1,141) and HPV− (n = 2,496) cells across the merged datasets. HPV− specimens show significantly higher IEG scores (median 1.828 vs. 1.452; Wilcoxon p = 2.74×10−17). *P < 0.05, **P < 0.01, ***P < 0.001. **l** Per-patient IEG+CD8_Tem proportions across both datasets. Proportion of IEG+CD8_Tem cells among total CD8+ T cells for each patient (GSM accession), stratified by dataset (GSE139324 vs. GSE164690) and HPV status. The subpopulation is consistently detectable across individual patients in both cohorts. **m** Proportion of FOS/JUN co-high-expressing (top 30%) CD8+ T cells by HPV status. Percentage of cells with combined IEG score ≥ 70th percentile (≥2.365) in HPV+ (24.3%, 277/1,141) versus HPV− (32.6%, 814/2,496) specimens. IEG+CD8_Tem cells constitute a substantial, reproducible fraction in both groups, confirming this fate-decision subpopulation as a conserved CD8+ T cell state independent of HPV etiology. Error bars, SEM; Wilcoxon rank-sum test.

