## Supplementary figures and images for "Integrated single-cell and spatial transcriptomic analysis of T cell exhaustion and immunometabolic remodeling in HPV-positive oropharyngeal squamous cell carcinoma"

### Figure S1

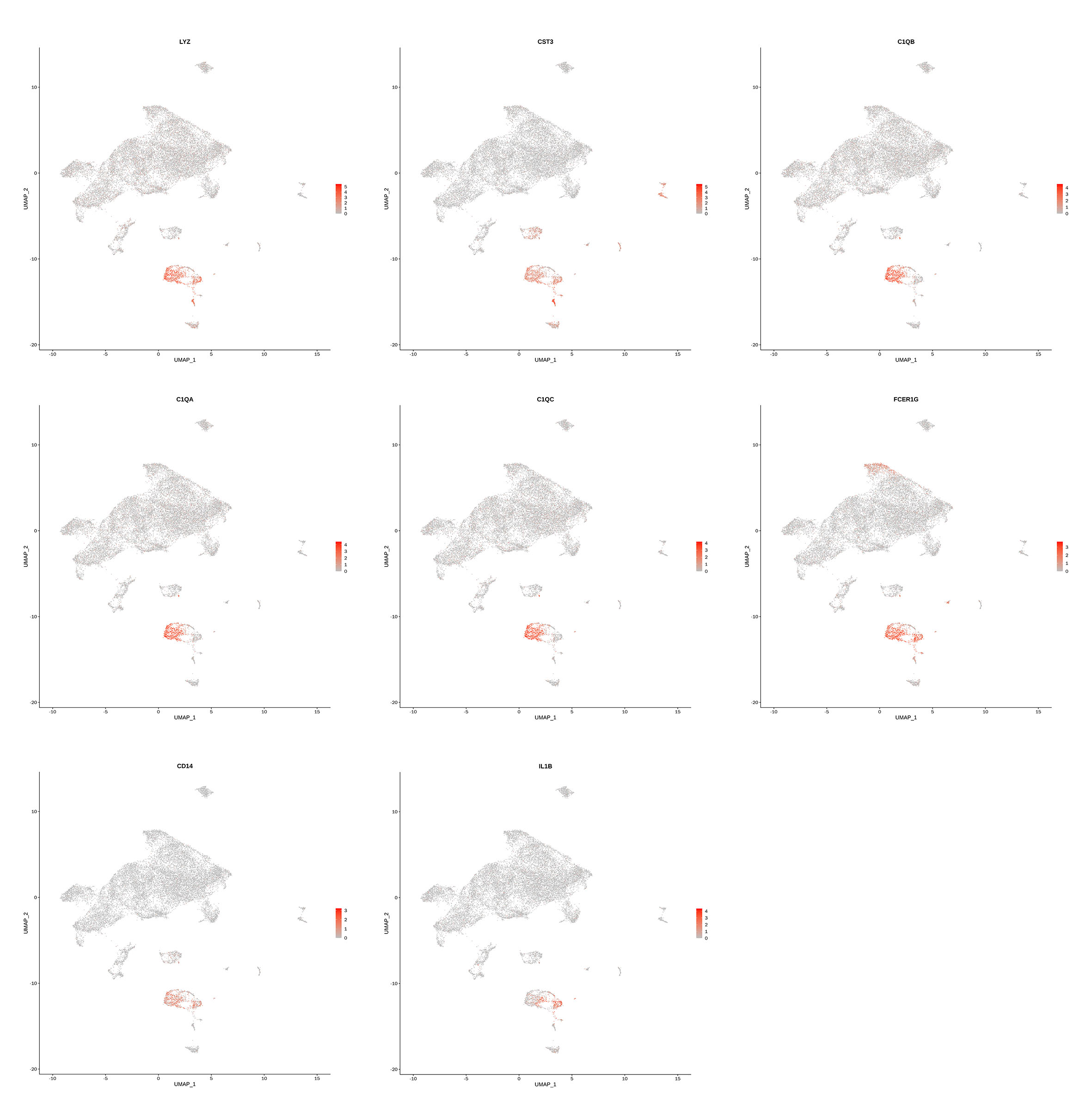

### Figure S2

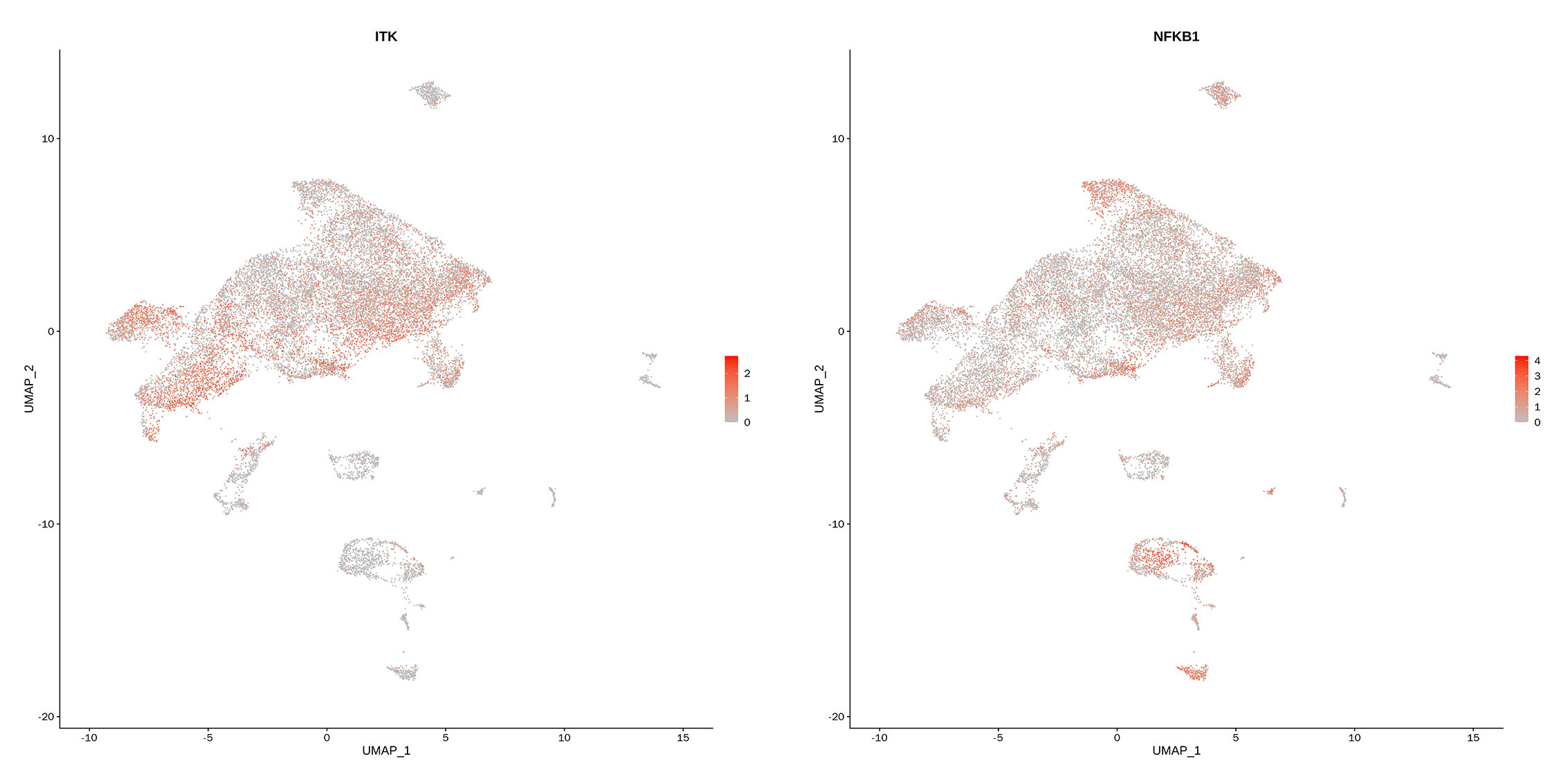

### Figure S3

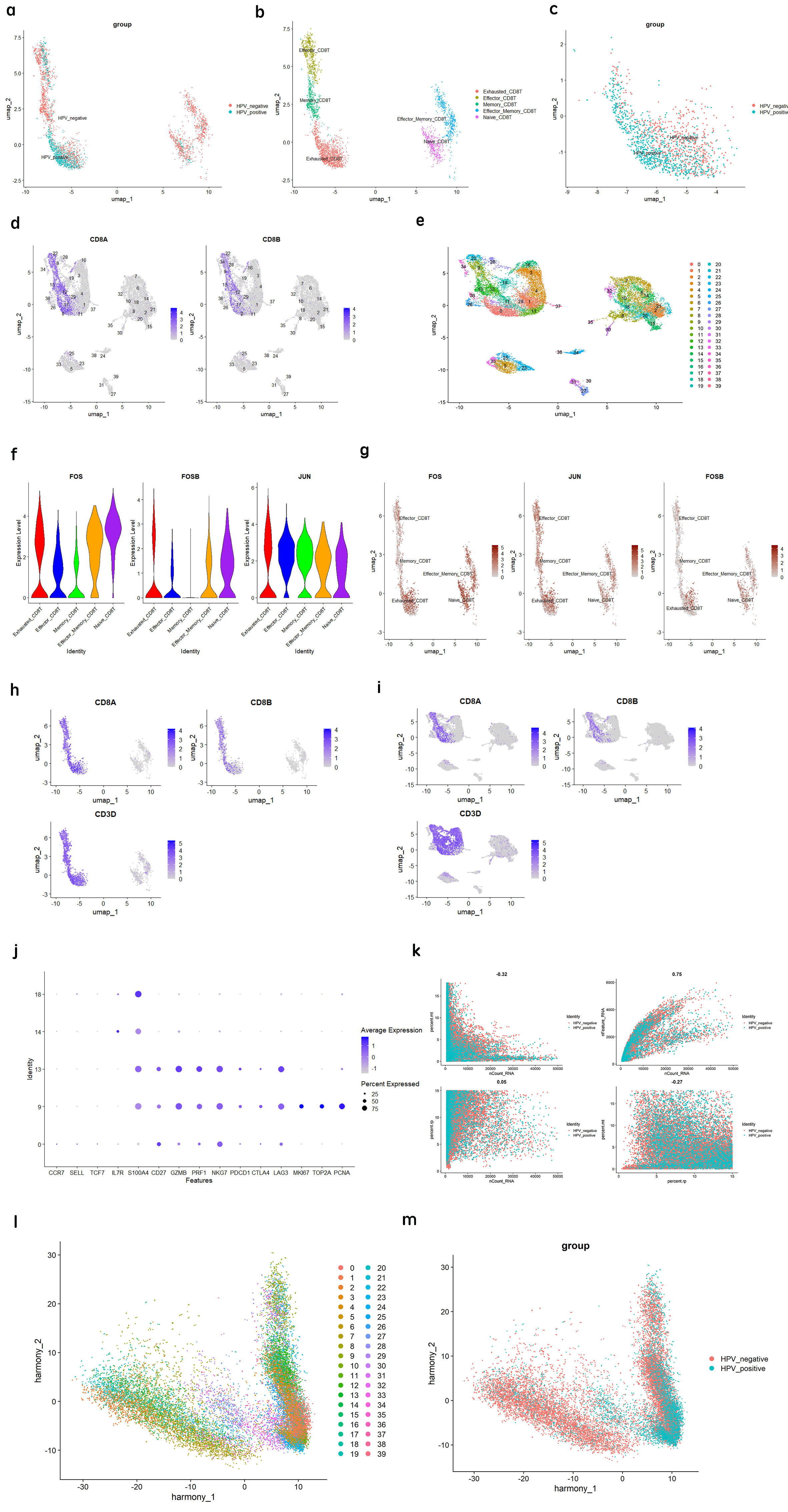

### Graphical Abstract

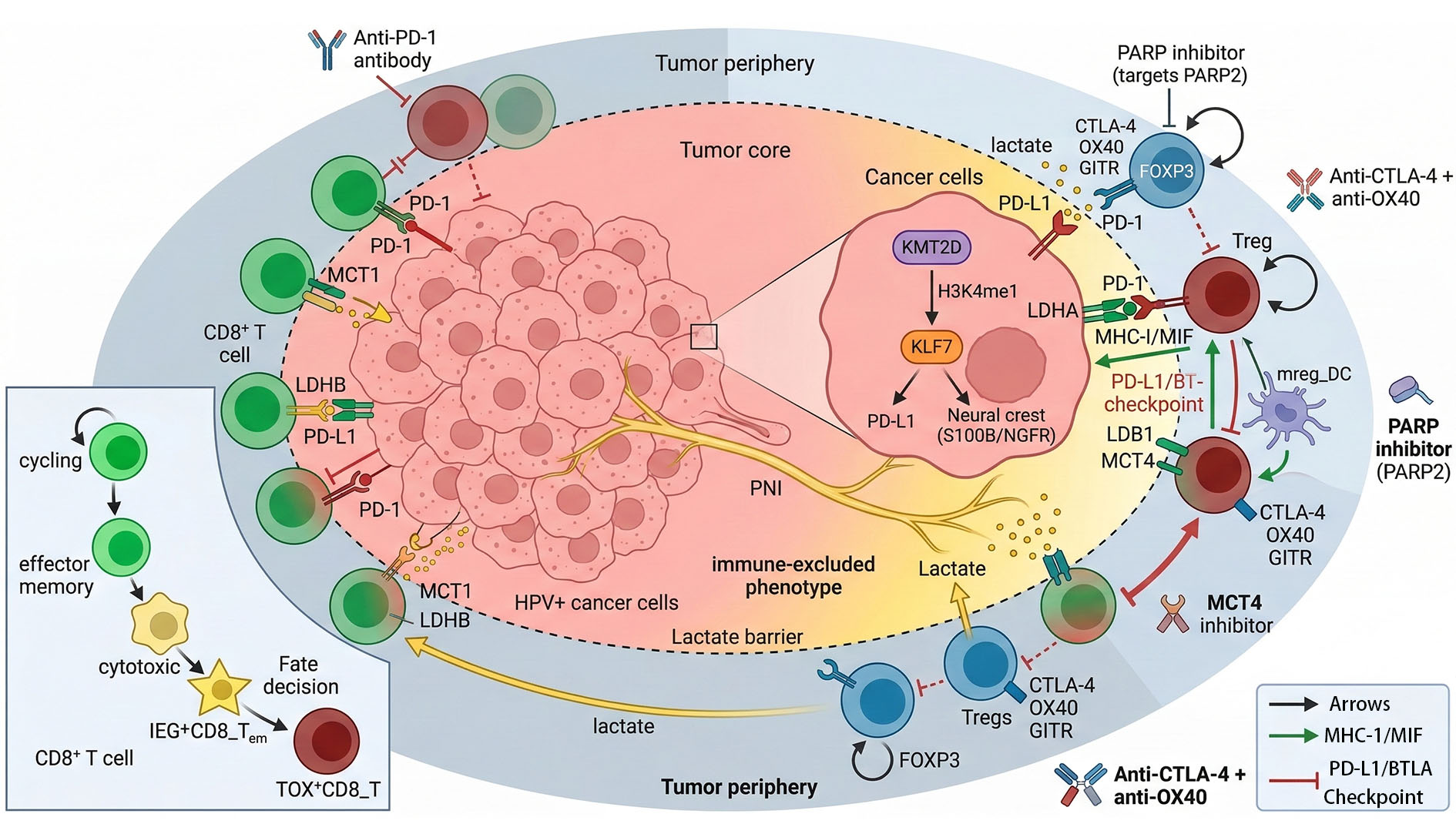
